# SARS-CoV-2 codon usage bias downregulates host expressed genes with similar codon usage

**DOI:** 10.1101/2020.05.05.079087

**Authors:** Andres Mariano Alonso, Luis Diambra

**Author notes:** Correspondence: Luis Diambra.

## Abstract

Severe acute respiratory syndrome is quickly spreading throughout the world and was declared as a pandemic by the World Health Organisation (WHO). The pathogenic agent is a new coronavirus (SARS-CoV-2) that infects pulmonary cells with great effectiveness. In this study we focus on the codon composition for the viral proteins synthesis and its relationship with the proteins synthesis of the host. Our analysis reveals that SARS-CoV-2 preferred codons have poor representation of G or C nucleotides in the third position, a characteristic which could conduct to an unbalance in the tRNAs pools of the infected cells with serious implications in host protein synthesis. By integrating this observation with proteomic data from infected cells, we observe a reduced translation rate of host proteins associated with highly expressed genes, and that they share the codon usage bias of the virus. The functional analysis of these genes suggests that this mechanism of epistasis contributes to understand some deleterious collateral effect as result of the viral replication. In this manner, our finding contribute to the understanding of the SARS-CoV-2 pathogeny and could be useful for the design of a vaccine based on the live attenuated strategy.

## 1 INTRODUCTION

The new SARS-CoV-2 coronavirus is the causative agent of the current pandemic of COVID-19. This highly pathogenic virus has quickly become the latest threat to the modern human lifestyle. Since the end of 2019 up to the redaction of this paper, this virus has infected over 3 million people, leading to mild symptoms from fever, lung function reduction, to severe acute respiratory syndrome (SARS) and even death. Today there are over 215.000 death all over the world. However, we are still far from determining the final mortality figure. In the absence of vaccines or effective antiviral treatments against SARS-CoV-2, it is important to understand how this virus appropriates the host translation apparatus and subverts the immune defences of infected cells. This can be the first step in the development of novel therapeutics.

As intracellular parasites, viruses replication mandatory depends on the translational machinery of their cellular hosts to translate viral transcripts. Thus, virus replication requires ribosomes, tRNA and translation factors from the cell host. On the other hand, codon usage bias is a feature subject to natural selection and affects the genomes from all domains of life. It is known that more frequently used codons are used for coding highly abundant proteins (30, 9, 35, 11). Virus genomes have also preference in the codon usage, but in this case the bias is constrained by the host translational machinery (38). The effects of codon composition of a transcript on its translation have been reported in literature (34, 13, 26, 39), and is considered an important determinant of gene expression (41, 42, 46). However, the codon usage of a gene also affects the translation of other genes (12). In fact the virus replication demands not only ribosomes, but also a lot tRNA resource for the most demanded codons. Thus, the consumption of specific tRNAs for the virus replication could be an alternative to control host protein synthesis machinery, as well as generating deleterious collateral effects on the function of the host cells.

Here, we study recent proteomic data from SARS-CoV-2 infected cells (4) from a novel point of view. By considering the codon usage of the virus ORFeome, we characterise a set of genes whose expression could be affected by the massive demand of the tRNA which implies the virus replication. We find that those host genes encoding proteins with similar codons to the virus ORFeome have lower translational rate. Extrapolating this finding to highly expressed genes in lung, we find a small set of genes that can be downregulated. These genes are involved in translation, immune systems, cell calcification, etc., and their roles on the SARS-CoV-2 cell pathogenesis should be the target of further studies.

## 2 RESULTS

We study codon composition of two coronavirus complete genomes by measuring the percentage of G and C nucleotides at the wobble position of the codons (GC3). Figure 1A depicts the GC3 content of each annotated coding sequence of the SARS-CoV and SARS-CoV-2. The comparison of the GC3 content reveals that codons preferentially used by the new virus have lower content of G and C at the third position than SARS-CoV. In particular, the ORFs corresponding to the proteins ORF1ab, ORF1a, surface, ORF6, ORF7a and ORF8 exhibit a very low content of GC3. In particular, the last two proteins are highly expressed at 10 hrs PI, as it can seen in Figure 1B.

**Figure 1.**
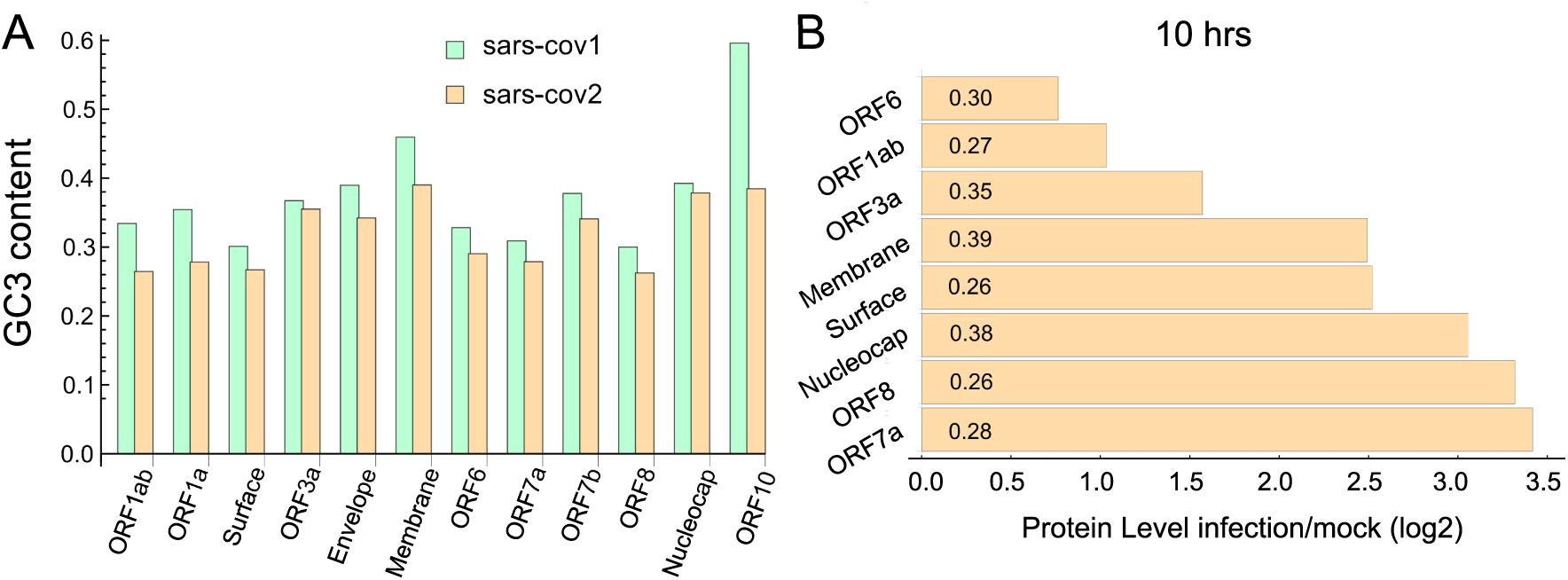
SARS-CoV-2 ORFome have lower content of GC3. **(A)** Bar chart comparing the G and C nucleotides on the third nucleotide (GC3) at every codon in the ORFeome of SARS-CoV and SARS-CoV-2. **(B)** Bar chart showing the proteins level of nine viral proteins at 10 hrs post infection (data obtained from (4)). The GC3 content for each viral ORF is showed in the corresponding bar.

Most of these proteins are highly expressed at 10h post infection (PI). This implies that the replication of SARS-CoV-2 would demand a higher supply for specific tRNAs than his ancestor. In turn, this could lead to an imbalance in the tRNA pool needed for the normal synthesis of the proteins of the host cell, altering its proteostasis. Of course, as tRNA pools depend on the cell types, the postulated imbalance could be a tissue-dependent feature. To check this hypothesis, we make use of the recently available data about the proteome profile and translational rate in SARS-CoV-2-infected cells (4). In this study CACO-2 cell line was infected with SARS-CoV-2 and mocked, the last one was used as control. From this proteome we select the coding sequences corresponding to the 100 most abundant proteins in the mock infected cells at 10h PI. Then, we compute the codon correlation (CCorr) between the codon usage of each one of these sequences and the codon usage of all coding sequences in SARS-CoV-2. Figure 2 depicts a raster plot of the translational rate of these genes from SARS-CoV-2-infected cells *versus* the corresponding CCorr (red dots). The black line is the adjusted linear model, which shows a significant and negative correlation between the translation rate and the codon composition of each sequence. This means that the translational rate of coding sequence in SARS-CoV-2-infected cells are lower than in mock-infected cells. Although the correlation (*ρ* = 0.26) with codon composition is not high, reflecting the fact that other factors could be regulating the translation rate, is significant at a of 0.05 (*p*-value= 0.024 *<* 0.05).

**Figure 2.**
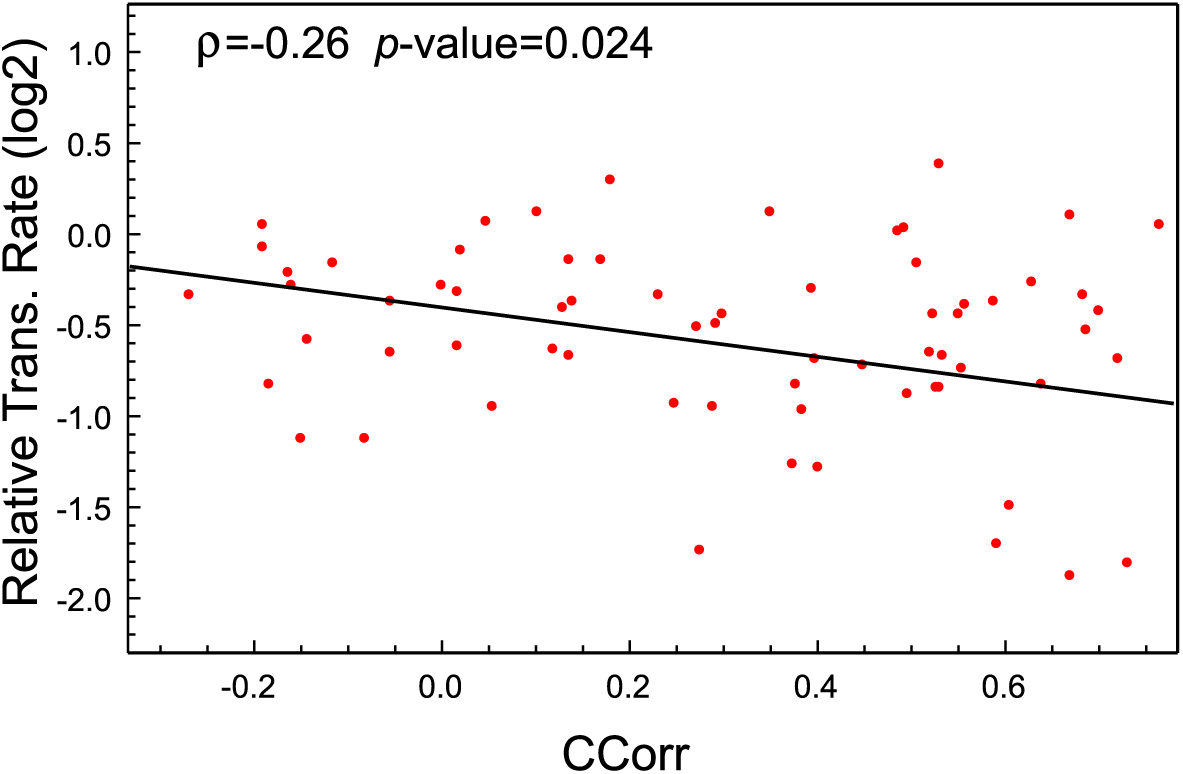
The translational rate of genes with high CCorr in SARS-CoV-2-infected cells is lower than the mock-infected cells. Raster plot showing negative correlation between translational rate of the 100 most expressed proteins in SARS-Cov-2 infected CACO-2 cells versus virus codon usage.

This analysis suggests then that the codon composition of highly expressed host-genes is a determinant of its own translation rate, and that viral replication induces a particular case of epistasis which could affect host cell proteostasis. Consequently, we find it relevant to extrapolate this phenomenon to the cells that conform lung tissue, the main target of the SARS-CoV-2 infection. To this aim, we select the 100 most expressed genes in lung from the GTEx database, and compute the CCorr for each sequence. Figure 3A shows that the codon composition profile of the most expressed genes from both cells types are different, it is evident that the lung cells share fewer codons in common with the SARS-CoV-2 than the CACO-2 cells. Consequently, the tRNAs pool used for the virus could be a more scarce supply in lung cells than in the CACO-2 cells.

**Figure 3.**
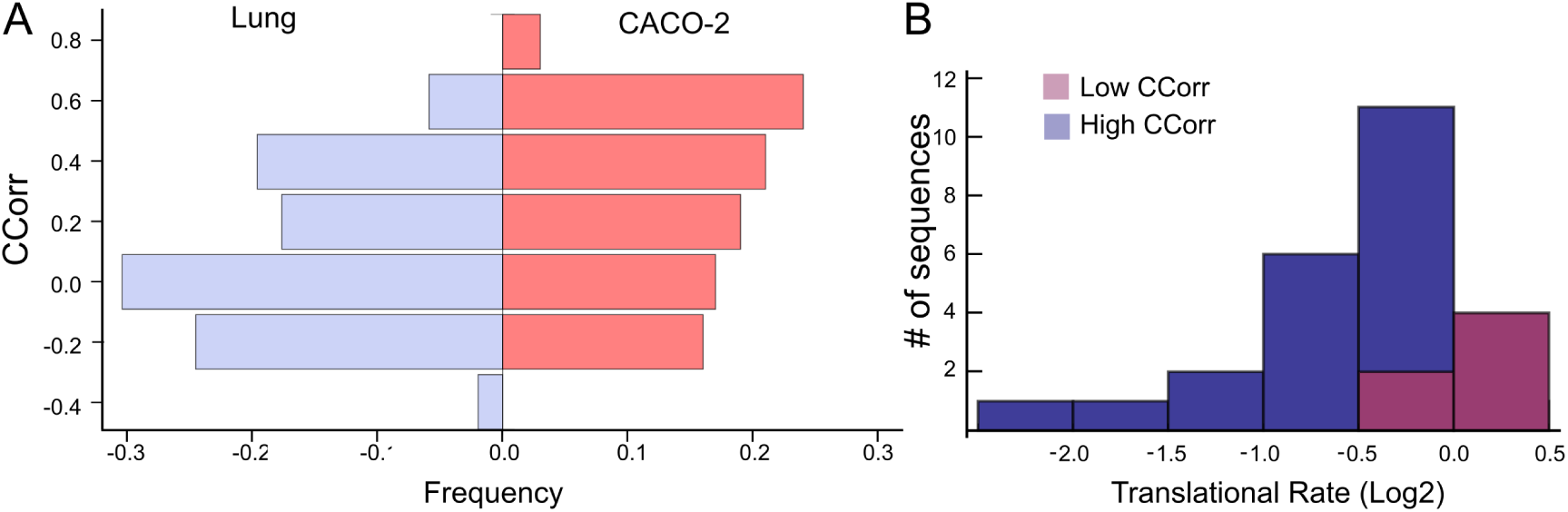
High codon correlation between SARS-CoV-2 and lung cells is associated with a translational rate decay over specific transcripts. **(A)** Frequency distribution of CCorr on the 100 highest expressed genes in lung cells and CACO-2 cells. **(B)** Histogram of the translational rate of 100 highest expressed genes in lung cells. Transcripts with high and low codon correlation (CCorr) with SARS-CoV-2 are highlighted.

We search for those genes, highly expressed in the lung tissue, which could be affected by the depletion of the tRNAs consumed by virus replication. They are those whose codon composition is similar to SARS-CoV-2 (CCorr *>* 0.25), they are listed in Table 1. Before analysing these 27 genes of interest, we compare the translation rate of these gene with the translation rate associated with highly expressed genes, but which differs in their codon usage bias (CCorr *<* −0.075). In this sense, we used the Mann-Whitney U test for median differences of independent samples to analyse the difference in translational rate between these two groups of genes. This test identified significant differences between these groups (*p*-value=0.0007), as it can be seen in Fig. 3B. We want to remark that, since there is still not available data on translational rate from lung cells exist, the last analysis was performed by using translational rate data available from CACO-2 cells, even knowing that this data would be underestimating the difference between these two groups of genes.

**Table 1.** Set of most expressed genes in lung (GTEx) and positive correlation with virus codon usage. col 1: gene name, col 2: CCorr, cols 4 and 5 translational rate, infected CACO-2 cells, measured at 6 and 10 hr PI, respectively. Translational rate are relative to mock cells.

| Gene Name | Codon corr | T. rate 6hs | T. rate 10hs |
| --- | --- | --- | --- |
| RPS3A | 0.582 | 0.686 | 0.980 |
| RPL5 | 0.559 | 0.585 | 0.271 |
| RPL9 | 0.546 | 0.723 | 0.738 |
| SPARCL1 | 0.536 | NA | NA |
| EEF1A1 | 0.531 | 0.948 | 0.627 |
| RPS12 | 0.443 | 0.155 | 0.649 |
| RPL17 | 0.438 | NA | 0.372 |
| TPT1 | 0.415 | 1.048 | 0.981 |
| RPL21 | 0.411 | NA | NA |
| RPS6 | 0.398 | 1.206 | 0.590 |
| RPL34 | 0.376 | 0.981 | 0.727 |
| RPS13 | 0.372 | 1.386 | 0.209 |
| RPL4 | 0.370 | 0.919 | 0.799 |
| RPS27A | 0.365 | NA | NA |
| RPSA | 0.352 | 0.902 | 0.643 |
| SAT1 | 0.350 | NA | NA |
| A2M | 0.340 | NA | NA |
| TXNIP | 0.317 | NA | NA |
| RPS25 | 0.315 | 0.728 | 0.753 |
| FN1 | 0.308 | NA | NA |
| RPL24 | 0.306 | 0.576 | 0.890 |
| RPLP1 | 0.303 | NA | NA |
| RPS7 | 0.297 | 0.965 | 1.036 |
| HSP90AB1 | 0.296 | 0.989 | 0.738 |
| B2M | 0.277 | 0.926 | 0.825 |
| S100A11 | 0.273 | 1.620 | 0.520 |
| MGP | 0.272 | NA | NA |

Further, we perform a GO term enrichment analysis over the 27 genes listed in Table 1, whose translation rate is decreased, by means of the Enrichr online software (7). Figure 4 illustrates the main enrichment pathways. This analysis reveals that codon usage could promote extensive changes in the translation machinery of the host in agreement with previous report in CACO-2 cells (4). Its is known that when canonical translation is impaired, as part of the host defence program, specific 40S ribosomal subunits are needed to support uncapped viral mRNA translation (22).

**Figure 4.**
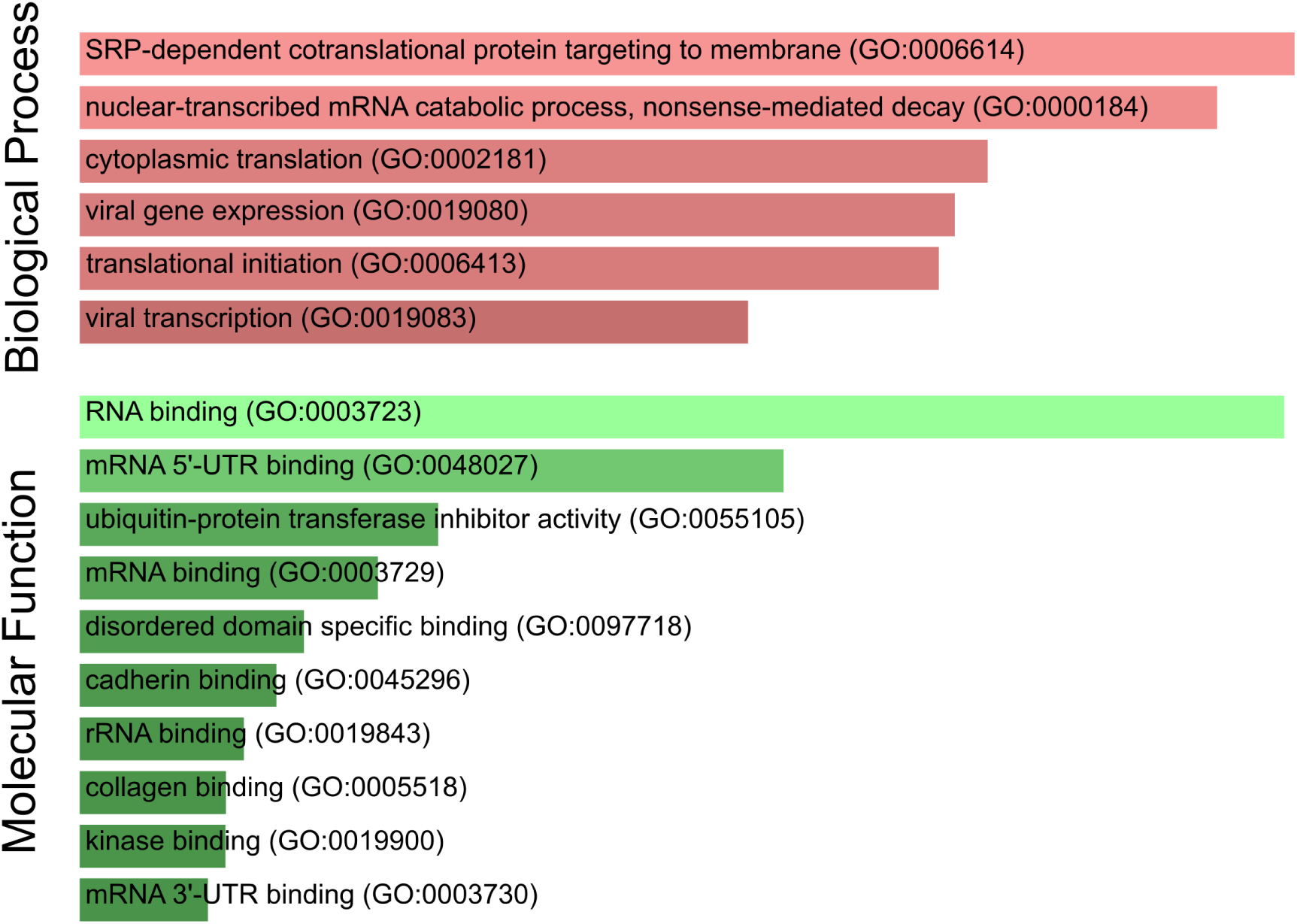
SARS-CoV-2 codon usage may have an impact on host translational machinery. GO enrichment analysis performed over genes of interest. The most enriched biological processes and molecular functions are highlighted.

Our list of genes is enriched with several ribosomal proteins that constitute both 40S (RPS6, RPS3A, RPS7, RPSA, RPS25, RPS13, RPS12, RPS27A) and 60S (RPL4, RPL5, RPL21, RPLP1, RPL34, RPL9, RPL24, RPL17) ribosomal subunits. All of them belong to the nonsense-mediated decay pathway that control the mRNAs with abnormal termination. Termination codon can be recognised if the 3’ untranslated region is short, or if it doess not have an exon junction complex downstream of the termination codon (28). These genes are also part of the process of SRP-dependent cotranslational protein destined for the endoplasmic reticulum.

It is known that these proteins are involved in translation, in particular the gene EEF1A1, an isoform of the alpha subunit of the elongation factor-1 expressed in lungs, which is responsible for the GTP-dependent binding of aminoacyl-tRNA to the ribosome. Beyond its function in translation, EEF1A1 takes part in the innate immune system, by activating directly the transcription of IFN-gamma (25). IFN-gamma activates immune cells, such as macrophages and natural killer cells and stimulates the major histocompatibility complex (MHC) II-dependent presentation of antigens to the immune system(37). This process could be downregulated due to a decrease in the translation rate of EEF1A1. Furthermore, B2M, other gene in the list of decreased translational rate, encodes one of the proteins that conform MHC class I found on the cell surface of all nucleated cells (2). Together with TXNIP, RPS3A and RPS27A, it is involved in the antigen processing and presentation. Moreover, this gene, along with S100A11, HSP90AB1 and also EEF1A1, regulates the exocytosis of granules containing inflammatory mediators in neutrophils (24).In addition, our analysis shows a translation decrease in A2M, a plasmatic protein that inhibits a broad spectrum of proteases, including trypsin, thrombin and collagenase (6). Finally, we highlight the presence in our analysis of the HSP90AB1, a known chaperone that facilitates the maturation of a wide range of proteins and its attenuation has been related to idiopathic pulmonary fibrosis and cystic fibrosis (16, 43).

In addition, we have also identified two genes (SAT1 and MGP) belonging to the pathway of endothelial cell calcification regulated by NOTCH1 (44). This finding acquire special relevance in the context of the acute lung injury (ALI) observed in many infected patients (18). The first gene SAT1 catalyses the acetylation of polyamines (spermidine and spermine) and carries it out of the cell. The polyamine excess is a prominent source of oxidative stress which can increases inflammatory response (20). Polyamines have also been connected with the immune system (32). On the other hand, the MGP gene encodes the matrix gla protein which is also highly expressed in all vasculatures. Recent studies suggest that MGP downregulates the tissue calcification by sequestering bone morphogenetic proteins (44). Mutations in this gene cause Keutel syndrome, which is characterised by peripheral pulmonary stenosis and abnormal cartilage calcification (27).

Other interesting protein in our study is SPARC-like 1 (SPARCL-1), also known as Hevin, commonly associated with regulation of cell migration, modulation of extracellular matrix proteins (15) and is has been shown to be involved in lymphocyte transendothelial migration through high endothelial venules (14). In this context, it is important to mention that clinical studies over patients that suffer several cases of COVID19 document a dysregulation of immune response related particularly with a lower lymphocytes count (18). In addition, our analysis reveals a translational rate decay of fibronectin (FN1), a master organiser of extracellular matrices that mediates cellular interactions playing important roles in cell adhesion, hemostasis and thrombosis (31, 43). According to our observation previously data show that SARS-CoV infected cells experiment a downregulation in fibronectin expression (40); as in other human virus like cytomegalovirus this last observation could be associated with pathogenesis of tissue inflammation (36).

## 3 DISCUSSION

The viral infection of human cells triggers an ensemble of host processes based on the interferons that interfere with viral replication. These processes have coevolved with the viral response to the host defence, thus virus counteract these process by a diversity of immuno-modulatory mechanisms. For example, NS1 protein plays a central role in the influenza infection by suppressing the host IFNs response. Further, the N protein of porcine reproductive and respiratory syndrome virus impairs the IFN transcription by acting over the TXK, which together with the EEF1A1 and PARP1 form the trimolecular complex that binds to the IFN-*γ* gene promoter (25, 21). Hemagglutinin of IAVs has been shown to facilitate IFNAR ubiquitination and degradation, reducing the levels of IFNAR, and thus suppressing the expression of IFN-stimulated antiviral proteins (45).

These examples illustrate the action of viral dedicated factors that downregulate the transcription of IFNs. Up to the present, no dedicated factor with analogue function have been identified in SARS-CoV and SARS-CoV-2. However, a recent report found a significant lack of IFN type I and III at the transcriptional level in human alveolar adenocarcinoma cells (3). On the other hand, a marked up-regulation of inflammatory mediators at the protein level (CXCL10, CCL2, IFN-*α* and *γ*) has been observed in patients with SARS-CoV, without a significant amount of specific antibody (5). Several case of absence of protective immunity due to previous infection seem to indicate a similar landscape for COVID-19. Until now, the manner in which some patients fail to develop adaptive immunity is yet to be elucidated. As mentioned in the Results section, the decreased translate of B2M could be related with this last observation since is a crucial factor for the stable presentation of antigens derived from virus or tumours proteins; this antigens are recognised by cytotoxic T cells that eventually eliminate the target cell stimulating apoptosis to prevent systemic dissemination of the disease (19).

MGP is also expressed at high levels in heart, kidney, and lung which is particular interesting in the context of several comorbidities and collateral effects observed in the COVID-19 patients (29). MGP is activated by a Vitamin K dependent carboxylase (10). Interestingly, an epidemiological study with hemodialysis patients showed that Vitamin K_2_ could prevent vascular calcification diseases (1). In contrast, the anticoagulant warfarin, which inhibits Vitamin K-dependent enzymes, was recently associated with an increased incidence of aortic valve calcification (17).

Summing up, if the depletion of a selected set of tRNA, induced by virus replication, affects the expression level or the co-translation folding of this proteins, one could expect the emergence of several systemic disorders.

## 4 CONCLUSION

Codon usage bias is thought to have significant effects on translation rate, where rare codons are assumed to be translated more slowly than common codons (33). It is assumed that rare and common codons are defined by usage rates of highly expressed genes. On the other hand, it is known that synonymous variations in a coding sequence can affects its folding and/or resulting expression level (26). However, whether the codon composition of viral ORFome can affect the translation rate of host genes has not been thoroughly explored. Here, we have shown that the synthesis of the proteins associated with highly expressed genes, and with similar codon usage to the one of the virus, appears to be downregulated. Following this idea we determine which genes in lung could be affected by the viral replication. A functional analysis of these genes reveal that they could be related to collateral effects observed in COVID-19 patients (18). Further studies are mandatory to corroborated or discard the putative relationship established here.

One of the main obstacles in the recent development of vaccines has been the finding of increased infectivity observed to occur after immunisations with whole virus vaccines or complete spike protein vaccines. This phenomenon has been observed both in vaccines against SARS coronavirus as in respiratory syncytial virus. On the other hand, other vaccine strategy has been recently assayed, focusing on altering the codon-pair usage without affecting protein sequence. This codon deoptimization strategy has been reduced virus replication (8, 23). We believe that our results shed light on how codon use could affect virus attenuation, help decrease the damaging side effect, providing an exciting opportunity for live-attenuated vaccine development.

## 5 MATERIALS AND METHODS

The coding sequences associated with the genome of SARS-CoV and SARS-CoV-2 were obtained from the NCBI (NC 004718.3 and NC 045512.2, respectively). Highly expressed genes in the lung tissue were retrieved from the GTEx portal (gtexportal.org). Proteomic and translatomic data from infected CACO-2 cells were retrieved from supplementary material of preprint (4), which are publicably available. These data contain protein levels from 6381 proteins and translational rate from 2715 proteins, consequently the proteomic information is not available for many proteins.

Given the coding sequence *s*, we compute the codon usage frequency as 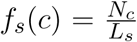, where *L*_*s*_ is total number of codons in the sequence *s*, and *N*_*c*_ is the number of times that codon *c* is present in *s*. In a similar manner, we define codon usage frequency relative to the viral ORFome: 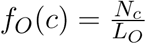, where *L*_*O*_ is the total number of codons in the ORFome, and *N*_*c*_ is the number of occurrences observed codon *c* in the ORFome. Thus, *f*_*s*_(*c*) and *f*_*O*_(*c*) are vector of 64 elements. To compute the CCorr of a given sequence *s* with viral ORFome we consider the Pearson’s correlation coefficient between the vector *f*_*s*_(*c*) and *f*_*O*_(*c*), as follows:

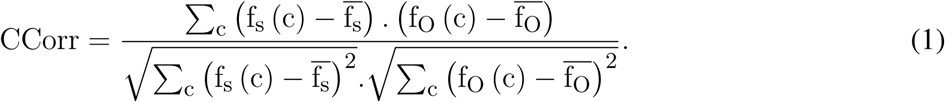

## ACKNOWLEDGMENTS

We would like to thank Nara Guisoni, Alejandra Carrea and Veronica Cóceres for their insightful suggestions and discussion. We are thankful to Catalina Trousdell for her judicious comments on the manuscript.

## DATA AVAILABILITY STATEMENT

The datasets analysed for this study can be found in the ProteomeXchange repository (ID=PXD017710) http://proteomecentral.proteomexchange.org/cgi/GetDataset?ID=PXD017710

